# The efficacy of obesity polygenic risk scores in an autistic population

**DOI:** 10.1101/836734

**Authors:** David Y. Zhang, Nathan P. Palmer, Chuan Hong, Luwan Zhang, Samuel G. Finlayson, Paul Avillach, Isaac S. Kohane

## Abstract

Polygenic risk scores (PRS) bear great promise in understanding complex diseases and improving clinical diagnoses, but the competency of these risk scores in different populations is known to vary significantly, especially between those of European ancestry and those of other ethnic ancestries. Additionally, the applicability of these risk scores across populations different by disease, instead of ethnicity, is poorly understood. A current and largely unexplored population for the accuracy of PRS is that of individuals with autism. Combined with the increased prevalence of obesity in autistic populations, we seek to evaluate the difference in efficacy of PRS for obesity in autistic versus non-autistic populations. We show that genetic variants strongly associated with obesity in non-autistic populations are significantly less representative of the disease in autistic populations. Rather, these cases of obesity phenocopies in patients with autism may follow a different and non-conventional mechanism of action involving the regulation of oxytocin in the brain among other potential behavioral factors. Our findings elucidate the limits of PRS across populations contrasting by disease and suggest that obesity may be regulated differently in individuals with autism as compared to those without autism.

## Introduction

Autism spectrum disorder (ASD) is a prevalent condition that affects a large proportion of the human population. Although it has been heavily studied in the past, there is still much more to investigate and learn about the disorder. One pattern that still remains unexplained is the increased prevalence of obesity in individuals with autism. Children and teenagers with autism are many times more likely to be overweight and to have obesity compared to healthy children [1, 2]. Furthermore, children with autism are at a higher risk for other related health issues including type 2 diabetes mellitus, hypertension, hyperlipidemia, and non-alcoholic fatty liver disease [3]. Without a clear genetic explanation, scientists have hypothesized behavioral reasons for these disparities. Children with autism are often picky and selective eaters which may increase susceptibility to unhealthy eating [4]. Additionally, dyspraxia in children with ASD may limit and inhibit physical activity [5]. These phenotypic manifestations might partially explain the increased prevalence of obesity in autistic populations, but we wanted to investigate this issue from a genetic perspective. Our study seeks to assess the effectiveness of using polygenic risk scores to identify obesity in patients with ASD in hopes of being able to intervene early and take preventative measures.

A polygenic risk score (PRS) takes advantage of the polygenic nature of complex traits and the small effects of common and low-frequency variants. In fact, common single nucleotide polymorphisms (SNPs) have been found to explain a large proportion of the heritability for complex traits [6, 7, 8, 9]. Using the sum of the effects of multiple common SNPs is often more telling than using a single rare SNP. Polygenic risk scores have been claimed to have great potential in predictive power for complex traits and henceforth important clinical utility as well [10]. However, there still remains much to learn about the generalizability of these risk scores across different traits as well as different populations.

In our study, we tested if the predictive power of a PRS can be applied in a population of individuals with autism for a complex trait such as obesity. There are already significant concerns regarding the use of polygenic risk scores across different ethnic populations [11, 12]. However, our study compares different populations, not by ethnicity, but by disease to observe if similar concerns arise. We first used the SNPs implicated in obesity from a non-autistic population to calculate obesity risk scores for children and young adults with autism from the Simons Simplex Collection. This evaluates how well the PRS performs in individuals with autism. Then we elucidated novel variants and pathways responsible for the increased prevalence of obesity in patients with ASD. We also evaluated the differences in explanatory power for obesity of single variants in non-autistic and autistic populations.

## Methods

The Simons Simplex Collection (SSC) used in our study is an aggregate of clinical and genetic data for over 2,500 autistic children between the ages of 4 and 18 [13]. Genetic information for parents and siblings is included as well. It is a resource provided by the Simons Foundation Autism Research Initiative (SFARI). For our study, single nucleotide polymorphisms were gathered for each of the 10,220 members of the Simons Simplex cohort, including patients and their families, using Illumina genotyping microarrays. Of the 10,220 samples, 1,354 were tested with the Illumina Human1M V1 C microarray, 4,626 with Illumina Human1M-Duo V3 microarray, and 4,240 with the Illumina HumanOmni 2.5-quad microarray. To facilitate comparisons across these data, the raw DNA sequences used in the probes from each microarray were mapped onto Human Reference genome GRCh38 using the BLAT alignment tool [14]. For sake of quality control, only probes that matched to a single location on the reference genome and with 100% identity were retained. The reference and alternative alleles used on the microarray probes were then annotated using the Ensembl Variant Effect Predictor (VEP) tool [15]. Measured values for autistic patient’s body mass index (BMI), age, and sex were used in our study as well.

A set of 20,088 genotyped patients from the Partners Biobank were included in our study as a non-autistic population control for our obesity risk scores. Patients are consented volunteers from seven medical institutions within the Partners Healthcare system. Samples were genotyped using three versions of the biobank SNP array offered by Illumina that is designed to capture the diversity of genetic backgrounds across the globe. The first batch of data (November 2015) was generated on the Multi-Ethnic Genotyping Array (MEGA) array, the first release of this SNP array. The second, third, and fourth batches (May 2016, Oct 2016, and Feb 2017) were generated on the Expanded Multi-Ethnic Genotyping Array (MEGA Ex) array. The fifth, sixth, and seventh batches (Feb 2017, Oct 2017) were generated on the Multi-Ethnic Global (MEG) BeadChip. MEGA was the initial release of a consortium array. As a consequence, a significant number of probes on this array were re-designed in later versions due to quality and probe re-synthesis issues. In order to ensure consistency, genotype data generated on MEGA and MEGA Ex were limited to the 1,416,020 SNPs and 1,741,376 SNPs, respectively, that appear on the final commercial release of this biobank array. All genetic data was mapped in reference to Human Reference genome GRCh38. Of the 20,088 individuals, BMI data was available for 18,100 patients. Measured values for each patient’s age, race, and sex were used in our study as well.

A set of 32 SNPs associated with obesity were used to calculate polygenic risk scores. These SNPs came from a study that used a systematic and replicable method for selecting the SNPs most strongly associated with obesity across multiple genome-wide association studies (GWAS) [16]. Furthermore, the use of this set of SNPs would allow for more valid comparisons between our results and the results from their study. Another set of 55 SNPs associated with obesity was compiled as well with the knowledge that a larger set of SNPs would help capture more of the variance for the phenotype. These SNPs came from two separate studies identifying genetic loci associated with body mass index [17, 18]. Tables of the two sets of SNPs and their weights can be found in the supplementary materials (**Supp. Table S1, S2**).

To calculate polygenic risk scores, we took a simple weighted sum of the numerical genotypes for the SNPs associated with obesity in a patient. Numerical genotypes were assigned as 0, 1, or 2 in reference to the effect allele used in calculating PRS. SNPs we used that were not present in a patient were imputed using the R package ‘snpStats’ [19]. Predictiveness of our risks scores was evaluated using three different metrics. Firstly, the Pearson’s correlation coefficient *r* was used to quantify the strength of the correlation between our calculated risk scores and BMI. Secondly, the area under the curve (AUC) for the receiver-operating characteristic (ROC) curve for obesity measures the probability that a randomly chosen obese patient will have a PRS greater than that of a randomly chosen non-obese patient. This metric served to evaluate the predictive power of our calculated risk scores. AUC calculations and ROC curves were generated using the R package ‘pROC’ [20]. Thirdly, a log odds-ratio was used to measure the strength of the association between obesity and our risk scores. For all statistical analyses, a p-value < 0.05 was defined as statistically significant. Obesity was classified as having a BMI≥30 and non-obese as BMI<30.

## Results

For all 2,317 sequenced autistic patients that we had BMI data for from the SSC, a polygenic risk score was computed using the set of 32 SNPs known to be associated with obesity (**Supp. Table S1**). Patients that were missing any of the 32 SNPs had those missing SNPs imputed prior to calculating the risk score. We then compared our calculated scores with the true BMI values for the cohort and found a weak and insignificant correlation. From **Figure 1A**, we see that the line of best-fit for a plot of BMI against obesity PRS has a near zero slope in this cohort of patients with ASD. We observe low risk scores for patients with a high BMI and high-risk scores for patients with a low BMI. To examine the correlation quantitatively, we correlated BMI and obesity PRS using the Pearson’s correlation coefficient and found *r* = 8.028E-03 (p-value = 0.699). We then calculated the same correlation coefficient for BMI and obesity PRS but controlling for demographic variables including age and sex and found *r* = 0.5761 (p-value = 2.2E-16). To compare our results with a baseline model, we also correlated BMI with only age and sex which gave *r* = 0.5764 (p-value = 2.2E-16).

**Figure 1.**
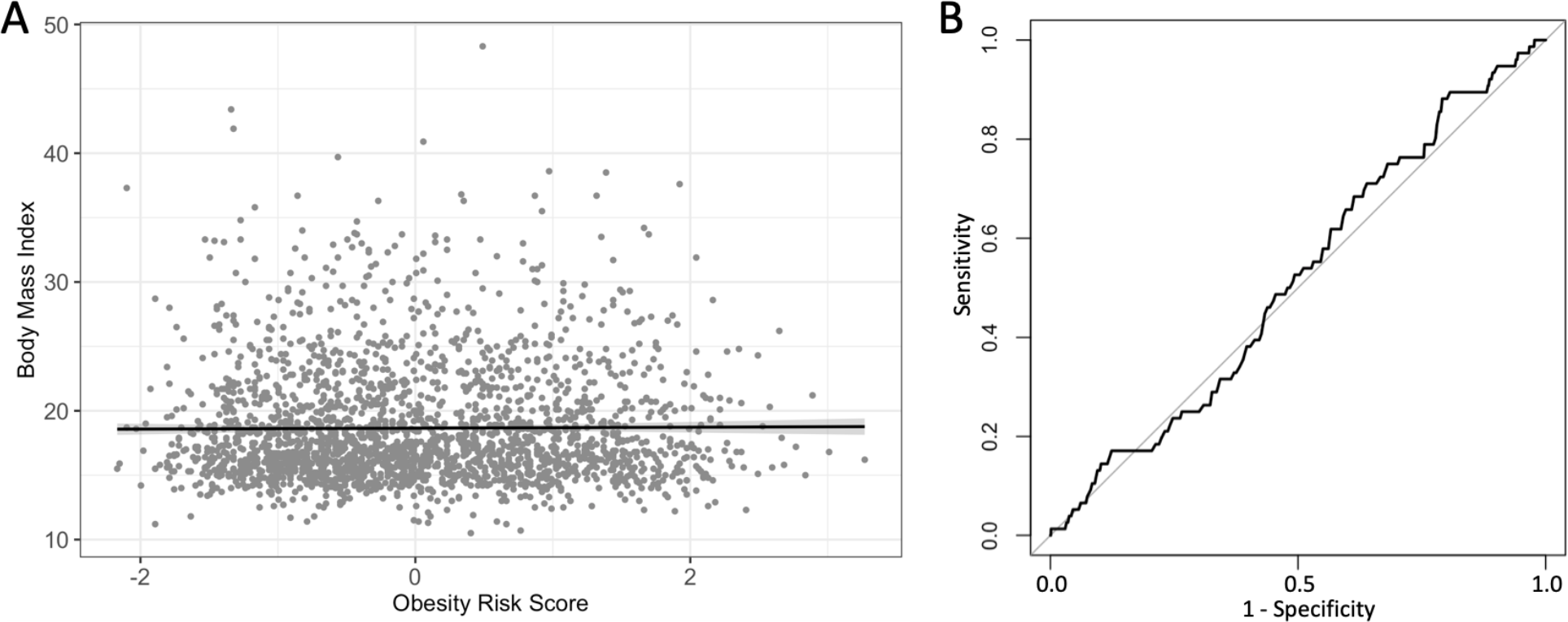
Plots generated using 2,317 sequenced autistic patients from the SSC and the set of 32 obesity-associated SNPs. (A) BMI measurements plotted against our calculated obesity polygenic risk scores. Scores on the x-axis are plotted as standard deviations from the mean. The line of best-fit represents a linear model of the data and has equation BMI~0.037(PRS) + 18.651. A 95% confidence interval for the linear model is shown as the gray shaded area around the line of best-fit. (B) A receiver-operating characteristic (ROC) curve for obesity (BMI≥30) using obesity PRS as the predictor.

Furthermore, we plotted a receiver-operating characteristic (ROC) curve to measure AUC, the area under the ROC curve, and evaluate the predictive power of our obesity risk scores in distinguishing between obese and non-obese. From **Figure 1B**, we see that our risk score is only marginally more sensitive than random choice (AUC = 0.516, 95% CI = 0.454-0.578).

To validate our methodology for computing obesity PRS, we replicated our calculations using a different cohort of non-autistic patients from the Partners Biobank (n = 18,100). Using the same set of 32 SNPs and identical methods for computing obesity PRS, we obtained risk scores for each patient in our new cohort. We then correlated our new scores with obesity using a logistic regression where obese was defined as BMI≥30. We also controlled for demographic factors including age, sex, and race. In this sample of non-autistic patients, our obesity risk score was positively and significantly correlated with BMI (*logOR* = 0.090, p-value = 0.002).

We then expanded our set of 32 SNPs to a set of 55 obesity associated SNPs (**Supp. Table S2**) in an attempt to capture more of the polygenic nature of obesity. Using the same Partners Biobank cohort of patients, new obesity risk scores were generated and correlated with obesity using a logistic regression while controlling for age, sex, and race. Compared to the regression computed using the original set of 32 SNPs in the same patient cohort, the new regression had a slightly larger and more significant log of odds ratio (*logOR* = 0.112 and p-value < 0.001).

Intrigued by the significance and strength of the results obtained using the Partner Biobank cohort, we used the new set of 55 obesity-associated SNPs to calculate new risk scores in the SSC cohort of patients with autism. Plotting BMI against these new risk scores produced a similar distribution as before when using the set of 32 SNPs, but **Figure 2A** exhibited a more evident positive correlation. In the new linear model for BMI and obesity PRS, the regression coefficient for obesity PRS was 0.250 whereas in the previous linear model generated using the set of 32 SNPs, the regression coefficient had been 0.037 (**Figure 1A**). The Pearson’s correlation coefficient was calculated for the new risk scores as well (*r* = 5.474E-02, p-value = 8.406E-03). When controlling for age and sex, *r* = 0.578 (p-value = 2.2E-16). A ROC curve was plotted to measure how well the new obesity risk scores predict obesity. Interestingly, the predictive power was poorer than before with a new AUC = 0.478 (95% CI = 0.416-0.539). This shows that for this cohort of autistic patients, obesity PRS could even be slightly worse than random guessing (**Figure 2B**).

**Figure 2.**
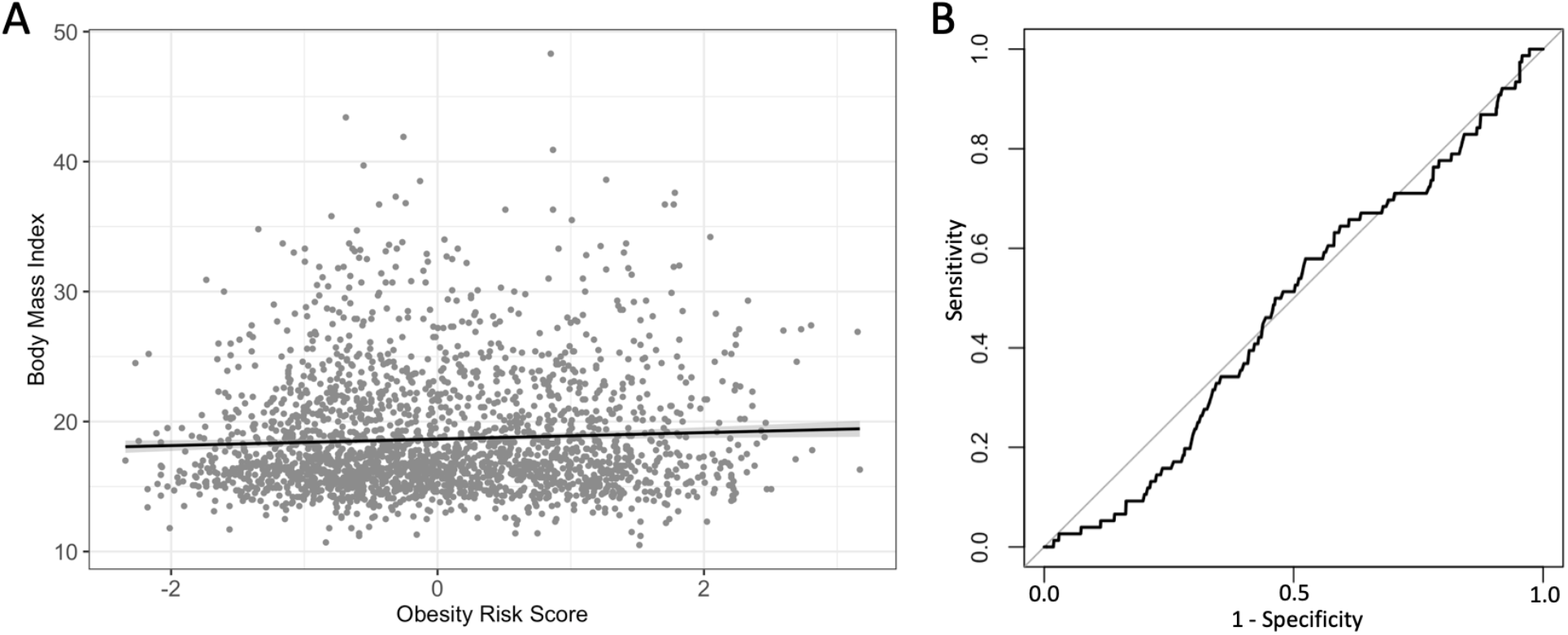
Plots generated using 2,317 sequenced autistic patients from the SSC and the set of 55 obesity-associated SNPs. (A) BMI measurements plotted against our calculated obesity polygenic risk scores. Scores on the x-axis are plotted as standard deviations from the mean. The line of best-fit represents a linear model of the data and has equation BMI~0.250(PRS) + 18.651. A 95% confidence interval for the linear model is shown as the gray shaded area around the line of best-fit. (B) A ROC curve for obesity (BMI≥30) using obesity PRS as the predictor.

Because the risk scores calculated from obesity-associated SNPs were weakly correlated with BMI in the SSC autistic cohort, we attempted to identify novel SNPs in the autistic patients associated with obesity. First, we performed single-SNP linear regressions for each of the 568,737 sequenced SNPs against BMI, controlling for age and sex. Treating each SNP independently does not account for interrelationships such as linkage disequilibrium but allows for the identification of all SNPs associated with BMI, either in a causal manner or not. Considering the high number of separate linear regressions being conducted, we applied a Bonferroni p-value correction to maintain a strict p-value cut-off for significance and to avoid the multiple comparisons problem. After p-value correction, only five SNPs had a p-value < 1.0 and only one of those SNPs had a statistically suggestive p-value < 0.1 (**Table 1**). With seemingly insignificant results, we ran a LASSO regression of BMI on a multivariate model including age, sex, and all 568,737 SNPs together. We ordered the input variables based on significance from our multiple single-SNP linear regressions to take advantage of LASSO’s iterative algorithm that converges when the model stabilizes and significantly reduces run-time. This regression uses shrinkage to eliminate variables unrelated to BMI and resulted in 6 SNPs with a non-zero regression coefficient (**Table 1**).

**Table 1.** The left side of the table contains all variables with a non-zero coefficient from our LASSO regression of BMI on age, sex, and all 568,737 SNPs. The right side of the table shows the results from multiple separate single-SNP linear regressions of BMI on age, sex, and each individual SNP. All SNPs from correlations with a p-value less than 1.0 after Bonferroni correction are shown below.

| LASSO regression |  | Multiple single-SNP linear regressions |  |  |
| --- | --- | --- | --- | --- |
| Variable | Coefficient | SNP | p-value | Coefficient |
| Intercept | 12.934 | rs12273646 | 0.097 | 19.488 |
| rs12273646 | 3.453 | rs11573755 | 0.184 | 19.016 |
| rs11573755 | 2.414 | rs7231745 | 0.358 | 9.285 |
| rs7231745 | 1.277 | rs37117 | 0.640 | 3.566 |
| age | 0.625 | rs9730633 | 0.871 | 3.464 |
| rs37117 | 0.433 |  |  |  |
| rs9730633 | 0.171 |  |  |  |
| rs909714 | 0.007 |  |  |  |

Observing that SNPs associated with obesity in the general population might not be indicative of obesity in an autistic cohort, we expanded our analysis to look for SNPs known to be associated with both BMI and autism. 941 SNPs that were known to be associated with BMI from the GIANT Consortium were compiled into a set [21] and 159 SNPs known to be associated with autism from The Autism Spectrum Disorders Working Group of The Psychiatric Genomics Consortium formed another set [22]. Comparing the two sets, only SNP rs4981693 was known to be associated with both phenotypes. The SNP is located within the non-coding RNA gene *LOC102724934*. We examined the expression of this SNP in obese versus non-obese autistic patients from our cohort. Of the 2,316 patients with autism who had genotype information on this SNP, 76 were obese (BMI≥30) and 2,240 were non-obese. An odds-ratio measuring the association of this SNP to obesity was calculated to be 1.160 with a 95% confidence interval of 0.692-1.675.

## Discussion

The results from using the initial 32 obesity-associated SNPs showed that the calculated risks scores were unindicative of BMI in an autistic population. The linear regression between BMI and risk score had a near-zero coefficient (**Figure 1A**) and the correlation was weak and insignificant. This was interesting because these exact same 32 SNP were used to calculate obesity risk scores in a non-autistic population done by another study. The weights for each SNP and the method used to calculate the risk score were identical [16]. In their study, they used the Atherosclerosis Risk in Communities cohort and found a correlation between BMI and obesity PRS of *r* = 0.13 (p-value < 1E-30) compared to our correlation coefficient of *r* = 8.028E-03 (p-value = 0.699). Furthermore, their risk score as a predictor for obesity had an AUC = 0.57 (95% CI 0.55-0.58) as opposed to our AUC = 0.516 (95% CI 0.454-0.578). This shows that the SNPs used as predictors for BMI and obesity in non-autistic populations are not similarly as useful in an autistic population. Our regression model between obesity and risk score using the Partner’s Biobank cohort also illustrated how these SNPs were more strongly and significantly correlated with obesity in a non-autistic population. This difference suggests that obesity phenocopies in individuals with autism might follow a different mechanism compared to cases of obesity in individuals without autism.

Our results from using an extended set of 55 obesity associated SNPs further supports our conclusion that obesity SNPs from a non-autistic population may not be representative of the phenotype in an autistic population. This larger set was an improved predictor for BMI in the SSC cohort as compared to the smaller set of 32 SNPs, but the correlation between BMI and risk score was still weak (*r* = 5.474E-02) and the AUC from using the risk score as a predictor of obesity (AUC = 0.478) fell below that of random choice. However, this set of SNPs still performed significantly better in the Partners Biobank cohort and was a much stronger predictor of obesity in a non-autistic population. The improved performance from a larger set of SNPs is to be expected as the additional SNPs can help explain more of the variance for a complex trait like BMI. This benefit has been noted and supported by other studies [23, 24]. A strong performance of risk scores as a predictor of BMI and obesity in non-autistic populations has been demonstrated using other populations as well such as the National Longitudinal Study of Adolescent Health Sibling Pairs [25]. In fact, the obesity SNPs that the authors used in that study largely overlap with the SNPs used in our study.

Although the reasons behind the significant difference in predictive power of obesity risk scores in our autistic cohort and other non-autistic cohorts is most likely due to pathway differences in the manifestation of obesity in these different populations, there could be other smaller contributing factors as well. One common and proven issue when it comes to comparing results between different populations is population stratification, the idea that different populations might have systematic differences in allele frequencies. Other studies have clearly outlined the presence of population stratification and its influences in association studies [11, 12], and this issue may be apparent in our study as well. Additionally, our autistic cohort from the Simons Simplex Collection is a population of autistic children from the ages of 4 to 18. The distribution of BMI has been shown to shift upwards as the age of a population increases [26] and given that the non-autistic cohorts used in our study generally consists of adults, discrepancies in the distribution of BMI between populations may have slightly affected our results as well. A comparison of the BMI distribution from our SSC cohort and from the general U.S. adult population can be found here (**Supp. Figure S1**).

The SNPs we found to be strongly correlated with BMI in our autistic cohort from running multiple single-SNP linear regressions were identical to the SNPs identified from our LASSO regression. These results give confidence that the identified SNPs play a role in obesity in autistic patients. Interestingly, none of the SNPs we identified have previously been implicated to be involved in or associated with obesity, possibly suggesting a novel obesity pathway specific to individuals with autism. However, plotting BMI against the genotypes of those SNPs in our cohort shows how our regression models might be overfitting to our sample (**Supp Figures 2A-F**). We are limited by the size of our autistic cohort and a larger sample size would be required to confirm the association of the variants with obesity. In addition, the SNP rs4981693 we found from other studies [21, 22] and known to be associated with both obesity and autism was not found to be significantly associated with obesity in our autistic cohort with an odds ratio that had a 95% confidence interval of 0.692-1.675. This suggests that this SNP does not offer the same explanatory power for obesity in patients with autism as opposed to those without autism, further supporting the prediction that another mechanism may be at play.

In trying to explain the increased prevalence of obesity in patients with autism, previous studies provide evidence of alternative pathways for the disease [1, 3]. One study has shown that subjects with autism have diminished grey matter in an area of the hypothalamus responsible for producing oxytocin [27]. Oxytocin is known to play a role in regulating feeding, specifically by reducing food intake and increasing energy expenditure [28, 29]. Therefore, perhaps a decrease in oxytocin levels in patients with autism may affect their ability to regulate eating that could lead to an increased risk for obesity. The hormone oxytocin has already been suspected of playing a role in the regulation of social behaviors in patients with autism [30]. With a growing body of evidence suggesting a role in other neuropsychiatric diseases as well including eating disorders, it is likely that the deregulation of the oxytocinergic system observed in autistic individuals provides a new pathway for obesity [31]. Additional studies will be required to learn more about the role of oxytocin in obesity for individuals with ASD and solidifying alternative pathways for the disease.

Our study found that well known SNPs associated with obesity are not as strongly representative of the disease in autistic populations. There are limitations to our study such as our lacking sample size that reduces the power of our regression models and prevents us from generalizing our results to larger populations. To further validate our results would require using a larger cohort of patients with ASD. In addition, more research in studying novel obesity pathways is needed, specifically those related to autism.

## Acknowledgments

We are thankful to all of the families at the participating Simons Simplex Collection (SSC) sites as well as the principal investigators at each site (A. Beaudet, R. Bernier, J. Constantino, E. Cook, E. Fombonne, D. Geschwind, R. Goin-Kochel, E. Hanson, D. Grice, A. Klin, D. Ledbetter, C. Lord, C. Martin, D. Martin, R. Maxim, J. Miles, O. Ousley, K. Pelphrey, B. Peterson, J. Piggot, C. Saulnier, M. State, W. Stone, J. Sutcliffe, C. Walsh, Z. Warren, E. Wijsman). We are grateful for obtaining access to genotypic and phenotypic data from the Simons Foundation Autism Research Initiative (SFARI) Base as well as from the Partners Healthcare Biobank. We also thank Chirag Patel for all the help and advice provided.

S.G.F. was supported by training grant T32GM007753 from the National Institute of General Medical Science.

## References

1. Broder-Fingert S., Brazauskas K., Lindgren K., Iannuzzi D. & Van Cleave J. Prevalence of Overweight and Obesity in a Large Clinical Sample of Children With Autism. Academic Pediatrics 14, 408–414 (2014).

2. Curtin C., Bandini L. G., Perrin E. C., Tybor D. J. & Must A. Prevalence of overweight in children and adolescents with attention deficit hyperactivity disorder and autism spectrum disorders: a chart review. BMC Pediatr 5, 48 (2005).

3. Shedlock K. et al. Autism Spectrum Disorders and Metabolic Complications of Obesity. The Journal of Pediatrics 178, 183–187.e1 (2016).

4. Cermak S. A., Curtin C. & Bandini L. G. Food selectivity and sensory sensitivity in children with autism spectrum disorders. J Am Diet Assoc 110, 238–246 (2010).

5. Dziuk M. A. et al. Dyspraxia in autism: association with motor, social, and communicative deficits. Developmental Medicine & Child Neurology 49, 734–739 (2007).

6. Yang J. et al. Common SNPs explain a large proportion of the heritability for human height. Nature Genetics 42, 565–569 (2010).

7. Zhu Z. et al. Dominance Genetic Variation Contributes Little to the Missing Heritability for Human Complex Traits. The American Journal of Human Genetics 96, 377–385 (2015).

8. Zhang Y., Qi G., Park J.-H. & Chatterjee N. Estimation of complex effect-size distributions using summary-level statistics from genome-wide association studies across 32 complex traits. Nature Genetics 50, 1318 (2018).

9. Speed D. et al. Reevaluation of SNP heritability in complex human traits. Nature Genetics 49, 986–992 (2017).

10. Torkamani A., Wineinger N. E. & Topol E. J. The personal and clinical utility of polygenic risk scores. Nature Reviews Genetics 19, 581 (2018).

11. Freedman M. L. et al. Assessing the impact of population stratification on genetic association studies. Nature Genetics 36, 388 (2004).

12. Hellwege J. et al. Population Stratification in Genetic Association Studies. Curr Protoc Hum Genet 95, 1.22.1–1.22.23 (2017).

13. Fischbach G. D. & Lord C. The Simons Simplex Collection: A Resource for Identification of Autism Genetic Risk Factors. Neuron 68, 192–195 (2010).

14. Kent W. J. BLAT—The BLAST-Like Alignment Tool. Genome Res. 12, 656–664 (2002).

15. McLaren W. et al. Deriving the consequences of genomic variants with the Ensembl API and SNP Effect Predictor. Bioinformatics 26, 2069–2070 (2010).

16. Belsky D. W. et al. Development and Evaluation of a Genetic Risk Score for Obesity. Biodemography and Social Biology 59, 85–100 (2013).

17. Speliotes E. K., Willer C. J., Berndt S. I., Monda K. L., Jackson A. U., Lango Allen H., … Loos R. J. F. Association analyses of 249,796 individuals reveal 18 new loci associated with body mass index. Nature Genetics 42, 937–948 (2010).

18. Hung C.-F., Breen G., Czamara D., Corre T., Wolf C., Kloiber S., … Rivera M. A genetic risk score combining 32 SNPs is associated with body mass index and improves obesity prediction in people with major depressive disorder. BMC Medicine 13, 86 (2015).

19. Clayton D. snpStats: SnpMatrix and XSnpMatrix classes and methods. R package version 1.34.0.

20. Robin X. et al. pROC: an open-source package for R and S+ to analyze and compare ROC curves. BMC Bioinformatics 12, 77 (2011)

21. Yengo L. et al. Meta-analysis of genome-wide association studies for height and body mass index in ∼700000 individuals of European ancestry. Hum Mol Genet 27, 3641–3649 (2018).

22. Anney R. J. L. et al. Meta-analysis of GWAS of over 16,000 individuals with autism spectrum disorder highlights a novel locus at 10q24.32 and a significant overlap with schizophrenia. Molecular Autism 8, 21 (2017).

23. Speakman J. R., Loos R. J. F., O’Rahilly S., Hirschhorn J. N. & Allison D. B. GWAS for BMI: a treasure trove of fundamental insights into the genetic basis of obesity. International Journal of Obesity 42, 1524 (2018).

24. Caballero A., Tenesa A. & Keightley P. D. The Nature of Genetic Variation for Complex Traits Revealed by GWAS and Regional Heritability Mapping Analyses. Genetics 201, 1601–1613 (2015).

25. Domingue B. W. et al. Polygenic Risk Predicts Obesity in Both White and Black Young Adults. PLOS ONE 9, e101596 (2014).

26. Flegal K. M. & Troiano R. P. Changes in the distribution of body mass index of adults and children in the US population. International Journal of Obesity 24, 807 (2000).

27. Kurth F. et al. Diminished Gray Matter Within the Hypothalamus in Autism Disorder: A Potential Link to Hormonal Effects? Biological Psychiatry 70, 278–282 (2011).

28. Noble E. E., Billington C. J., Kotz C. M. & Wang C. Oxytocin in the ventromedial hypothalamic nucleus reduces feeding and acutely increases energy expenditure. American Journal of Physiology-Regulatory, Integrative and Comparative Physiology 307, R737–R745 (2014).

29. Sabatier N., Leng G. & Menzies J. Oxytocin, feeding, and satiety. Front Endocrinol (Lausanne) 4, 35 (2013).

30. Andari E. et al. Promoting social behavior with oxytocin in high-functioning autism spectrum disorders. Proc Natl Acad Sci U S A 107, 4389–4394 (2010).

31. Romano A., Tempesta B., Micioni Di Bonaventura M. V. & Gaetani S. From Autism to Eating Disorders and More: The Role of Oxytocin in Neuropsychiatric Disorders. Front Neurosci 9, (2016).

32. Schwartz L. M. & Woloshin S. Changing disease definitions: implications for disease prevalence. Analysis of the Third National Health and Nutrition Examination Survey, 1988-1994. ECP 2, 76–85 (1999).

