## Supplementals for "The efficacy of obesity polygenic risk scores in an autistic population"

**Table S1**. The set of 32 SNPs identified to be strongly associated with obesity and used to calculate polygenic risk scores in the Simons Simplex Collection and the Partners Biobank. These SNPs were collected from this study [16].

| SNP_ID | chr_location | effect_allele | weight |
|---|---|---|---|
| rs2815752 | 1_72346757 | G | 0.13 |
| rs1514175 | 1_74525960 | A | 0.07 |
| rs1555543 | 1_96479241 | A | 0.06 |
| rs543874 | 1_177920345 | G | 0.22 |
| rs759250 | 2_59102018 | A | 0.1 |
| rs2121279 | 2_142285716 | T | 0.08 |
| rs2867123 | 2_624524 | G | 0.3 |
| rs10182181 | 2_24927427 | G | 0.14 |
| rs12714640 | 3_85800912 | A | 0.1 |
| rs1516728 | 3_186112102 | T | 0.11 |
| rs12641981 | 4_45177866 | T | 0.18 |
| rs13114738 | 4_102363708 | T | 0.13 |
| rs10057967 | 5_75701931 | C | 0.1 |
| rs6864049 | 5_124994829 | A | 0.07 |
| rs734597 | 6_50868566 | A | 0.13 |
| rs1412235 | 9_28410998 | C | 0.11 |
| rs867559 | 9_126703046 | G | 0.24 |
| rs2028882 | 11_8483307 | C | 0.06 |
| rs10501087 | 11_27648561 | C | 0.18 |
| rs12419692 | 11_47603162 | A | 0.05 |
| rs7138803 | 12_49853685 | A | 0.12 |
| rs1475219 | 13_27463278 | C | 0.09 |
| rs1440983 | 14_30024055 | A | 0.15 |
| rs7144011 | 14_79474040 | T | 0.13 |
| rs28670272 | 15_67817569 | G | 0.13 |
| rs11639988 | 16_19933041 | G | 0.17 |
| rs12443881 | 16_28830456 | T | 0.15 |
| rs9939609 | 16_53786615 | A | 0.38 |
| rs12970134 | 18_60217517 | A | 0.21 |
| rs11084753 | 19_33831232 | A | 0.04 |
| rs11083779 | 19_45701620 | C | 0.07 |
| rs7250850 | 19_47082260 | G | 0.09 |

**Table S2**. The set of 55 SNPs identified to be strongly associated with obesity and used to calculate polygenic risk scores in the Simons Simplex Collection and the Partners Biobank. These SNPs were collected from these studies [17, 18].

| SNP_ID | chr_location | effect_allele | weight |
|---|---|---|---|
| rs1558902 | 16_53769662 | A | 0.39 |
| rs2867125 | 2_622827 | C | 0.31 |
| rs571312 | 18_60172536 | A | 0.23 |
| rs10938397 | 4_45180510 | G | 0.18 |
| rs10767664 | 11_27704439 | A | 0.19 |
| rs2815752 | 1_72346757 | A | 0.13 |
| rs7359397 | 16_28874338 | T | 0.15 |
| rs9816226 | 3_186116710 | T | 0.14 |
| rs3817334 | 11_47629441 | T | 0.06 |
| rs29941 | 19_33818627 | G | 0.06 |
| rs543874 | 1_177920345 | G | 0.22 |
| rs987237 | 6_50835337 | G | 0.13 |
| rs7138803 | 12_49853685 | A | 0.12 |
| rs10150332 | 14_79470621 | C | 0.13 |
| rs713586 | 2_24935139 | C | 0.14 |
| rs12444979 | 16_19922278 | C | 0.17 |
| rs2241423 | 15_67794500 | G | 0.13 |
| rs2287019 | 19_45698914 | C | 0.15 |
| rs1514175 | 1_74525960 | A | 0.07 |
| rs13107325 | 4_102267552 | T | 0.19 |
| rs2112347 | 5_75719417 | T | 0.1 |
| rs10968576 | 9_28414341 | G | 0.11 |
| rs3810291 | 19_47065746 | A | 0.09 |
| rs887912 | 2_59075742 | T | 0.1 |
| rs13078807 | 3_85835000 | G | 0.1 |
| rs11847697 | 14_30045906 | T | 0.17 |
| rs2890652 | 2_142202362 | C | 0.09 |
| rs1555543 | 1_96479241 | C | 0.06 |
| rs4771122 | 13_27446043 | G | 0.09 |
| rs4836133 | 5_124996410 | A | 0.07 |
| rs4929949 | 11_8583046 | C | 0.06 |
| rs206936 | 6_34335092 | G | 0.06 |
| rs2568958 | 1_72299433 | A | 0.13 |
| rs11165643 | 1_96458541 | T | 0.06 |
| rs10913469 | 1_177944384 | C | 0.22 |
| rs10182181 | 2_24927427 | G | 0.14 |

| | | | |
|---|---|---|---|
| rs759250 | 2_59102018 | A | 0.1 |
| rs6714473 | 2_142283513 | T | 0.09 |
| rs7640855 | 3_85825014 | A | 0.1 |
| rs7647305 | 3_186116501 | C | 0.14 |
| rs12641981 | 4_45177866 | T | 0.18 |
| rs253414 | 5_75660692 | T | 0.1 |
| rs6864049 | 5_124994829 | A | 0.07 |
| rs2183825 | 9_28412377 | C | 0.11 |
| rs10840065 | 11_8505096 | A | 0.06 |
| rs6265 | 11_27658369 | C | 0.19 |
| rs10838738 | 11_47641497 | G | 0.06 |
| rs1475219 | 13_27463278 | C | 0.09 |
| rs10146997 | 14_79478819 | G | 0.13 |
| rs12446632 | 16_19924067 | G | 0.17 |
| rs4788102 | 16_28862077 | A | 0.15 |
| rs3751812 | 16_53784548 | T | 0.39 |
| rs921971 | 18_60194430 | C | 0.23 |
| rs2303108 | 19_47086638 | C | 0.09 |
| rs11083779 | 19_45701620 | T | 0.15 |

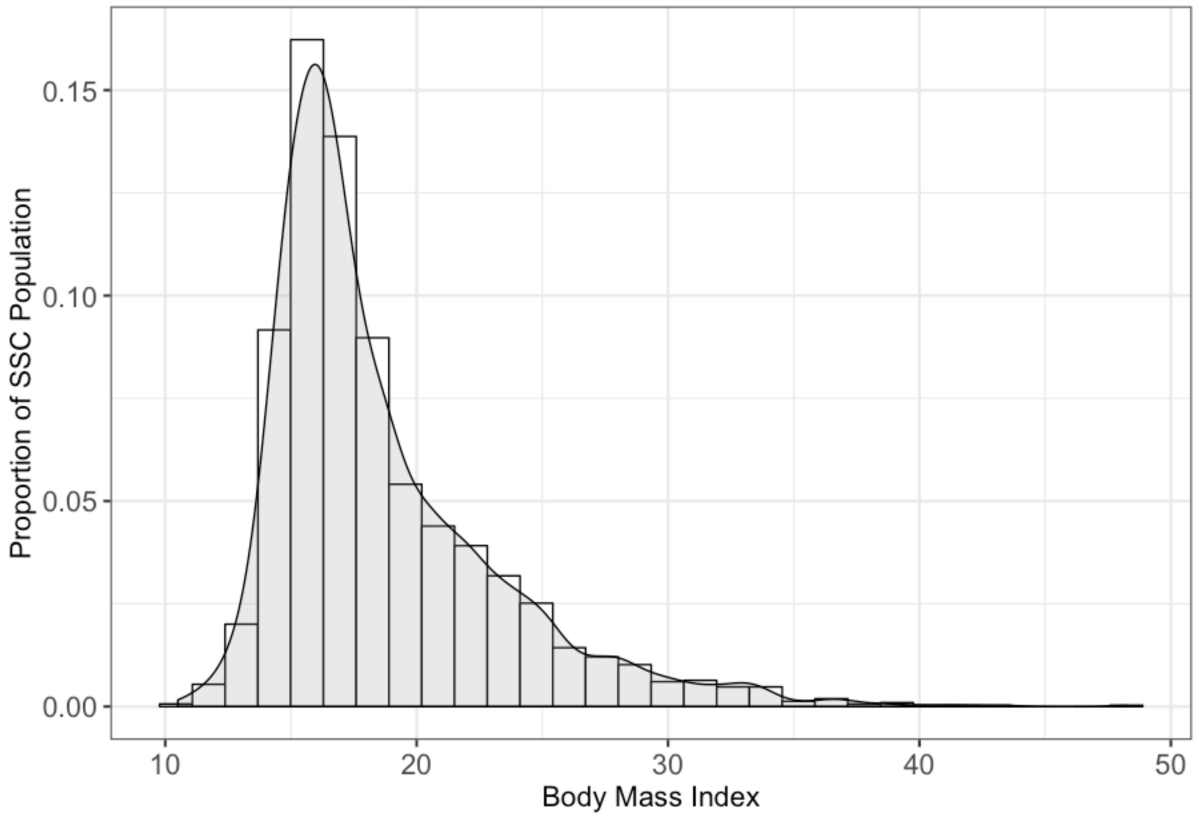

**Figure S1**. A BMI distribution of the 2,317 patients from the Simons Simplex Collection. This is compared to the distribution of BMI for the U.S. adult population in Figure 5 of this study [32].

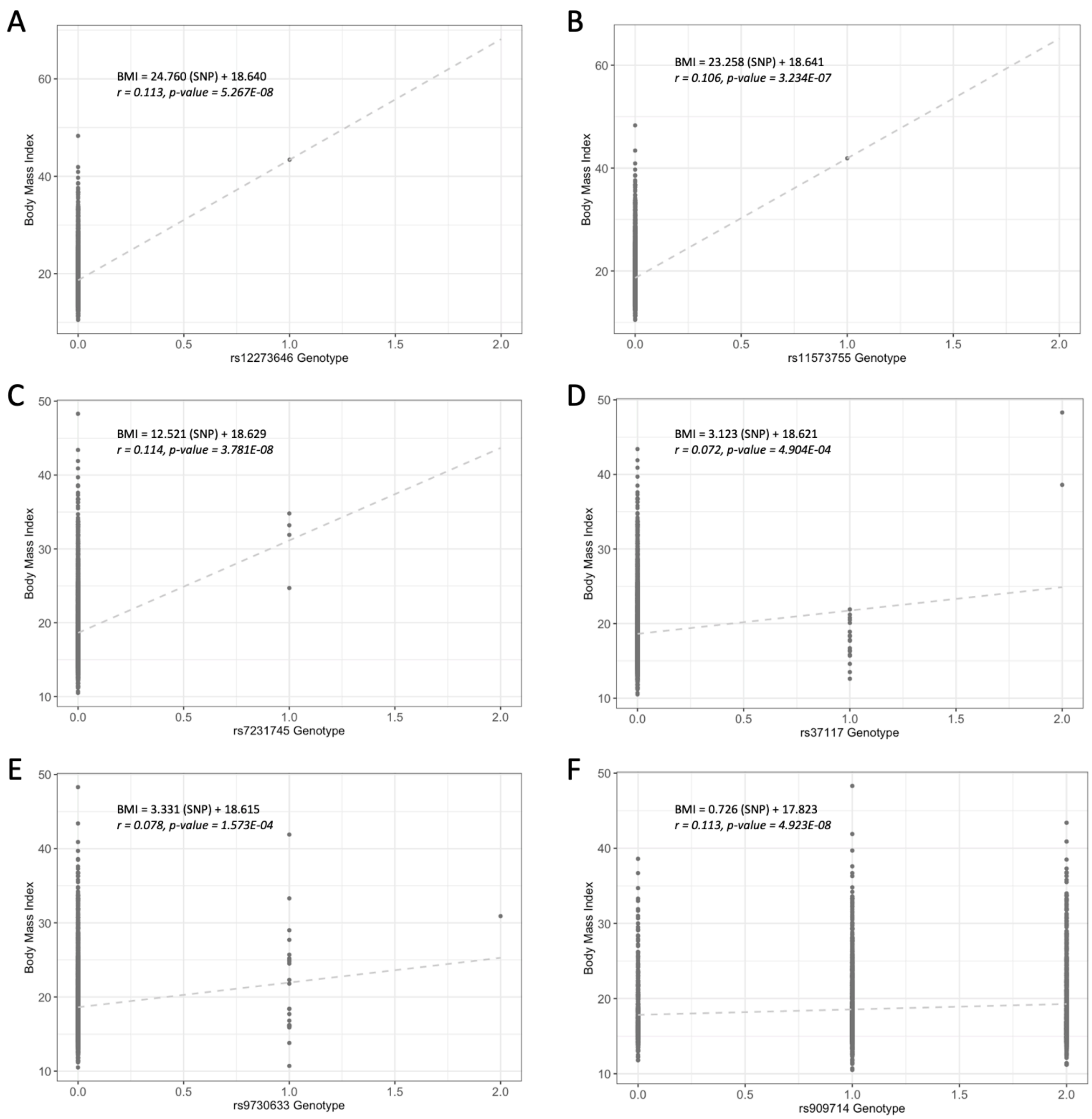


**Figure S2**. (A-F) Plots of BMI against the numerical genotypes of the six SNPs identified as being associated with obesity in our SSC cohort by either multiple single-SNP linear regressions or LASSO regression. The equation for the linear model of each regression is displayed for each plot as well as the Pearson's correlation coefficient and its p-value. These plots suggest potential overfitting of our models and will require a larger sample size to confirm our results.
